# HumanTestisDB: A Comprehensive Atlas of Testicular Transcriptomics and Cellular Interactions

**DOI:** 10.1101/2024.11.06.622208

**Authors:** Mengjie Wang, Laihua Li, Qing Cheng, Hao Zhang, Zhaode Liu, Yiqiang Cui, Jiahao Sha, Yan Yuan

## Abstract

Advances in single-cell technology have enabled the detailed mapping of testicular cell transcriptomes, which is essential for understanding spermatogenesis. However, the fragmented nature of age-specific data from various literature sources has hindered comprehensive analysis. To overcome this, the Human Testis Database (HumanTestisDB) was developed, consolidating multiple human testicular sequencing datasets to address this limitation. Through extensive investigation, 38 unique cell types were identified, providing a detailed perspective on cellular variety. Furthermore, the database systematically categorizes samples into eight developmental stages, offering a structured framework to comprehend the temporal dynamics of testicular development. Each stage features comprehensive maps of cell-cell interactions, elucidating the complex communication network inside the testicular microenvironment at particular developmental stages. Moreover, by facilitating comparisons of interactions among various cell types at different stages, the database permits examining alterations that transpire during critical transitions in spermatogenesis. HumanTestisDB, available at https://shalab.njmu.edu.cn/humantestisdb, offers vital insights into testicular transcriptomics and interactions, serving as an essential resource for advancing research in reproductive biology.

## Introduction

In the male reproductive system, the human testis is essential for both sperm production and hormone regulation. It includes various cell types, all essential to the spermatogenesis process [1–3]. Single-cell technologies have advanced significantly in recent years, offering previously unheard-of insights into the testicular microenvironment’s functional dynamics and cellular heterogeneity [4–11]. These technologies have been applied to human fetal germ cells (FGCs), demonstrating that the development of male FGCs undergoes stages including migration, mitosis, and cell-cycle arrest [4]. The transcriptome profiles of adult testes were then generated, describing the different states of spermatogonia and analyzing gene expression changes throughout male meiosis and spermiogenesis [5–7]. Following constructing the transcriptome maps of the testes during embryonic, fetal, childhood, and adolescence, further understanding of the development of somatic cells, particularly Sertoli and Leydig cells, was achieved [8–11]. Despite these developments, the testicular sequencing data remains unintegrated and fails to illustrate the distribution of cell types across various ages. Furthermore, alternative analytical methodologies employed in diverse literature yield incongruent results.

This study tackles the significant deficiency in consolidating and analyzing disparate testicular cell data. Our contribution, HumanTestisDB, signifies a substantial advancement in the compilation and augmentation of existing testicular cell datasets with comprehensive annotations. The purpose of this database is to function as a repository and to offer a framework for comprehending the complex cell-cell interactions within the testicular microenvironment at different developmental stages. HumanTestisDB provides essential resources for researchers and clinicians to investigate the intricate cellular processes and molecular pathways underlying testicular development and function. Infertility impacts a large number of couples globally, with male factor infertility constituting a substantial share [12, 13]. However, the primary reasons for male infertility, including azoospermia (the lack of sperm in semen), remain inadequately comprehended [14]. Researchers can better understand the molecular basis of male infertility by comparing the HumanTestisDB data with sequencing data from azoospermia patients. This understanding could inform future research, encourage individualized medical approaches to infertility treatment, and eventually improve the reproductive outcomes of those who are affected.

Multiple human testicular sequencing datasets covering a broad age range were methodically combined to develop HumanTestisDB. This study revealed 38 different cell types, providing a thorough understanding of the cellular makeup of the testis. After classifying samples from different ages into eight stages, the complex cellular makeup and interactions inside each stage were analyzed. The molecular mechanisms underlying testicular development and function can be better understood using this stratification to uncover universal and stage-specific signaling pathways.

## Results

### Overview of HumanTestisDB

HumanTestisDB is a comprehensive database that includes 46 single-cell sequencing datasets from human testis, ranging from 6 weeks post-fertilization (W6) to 49 years of age (Y49) (**Figure 1**A, **Figure 2**B, Table S1). Following quality control, our final dataset comprised 131,113 cells (Figure S1). Employing established marker expression patterns (Figure 2C, D, Figure S2), these cells were initially classified into 18 distinct cell types (Figure 2A). This classification comprised 13 somatic and five germ cell types (Figure 2A).

**Figure 1.**
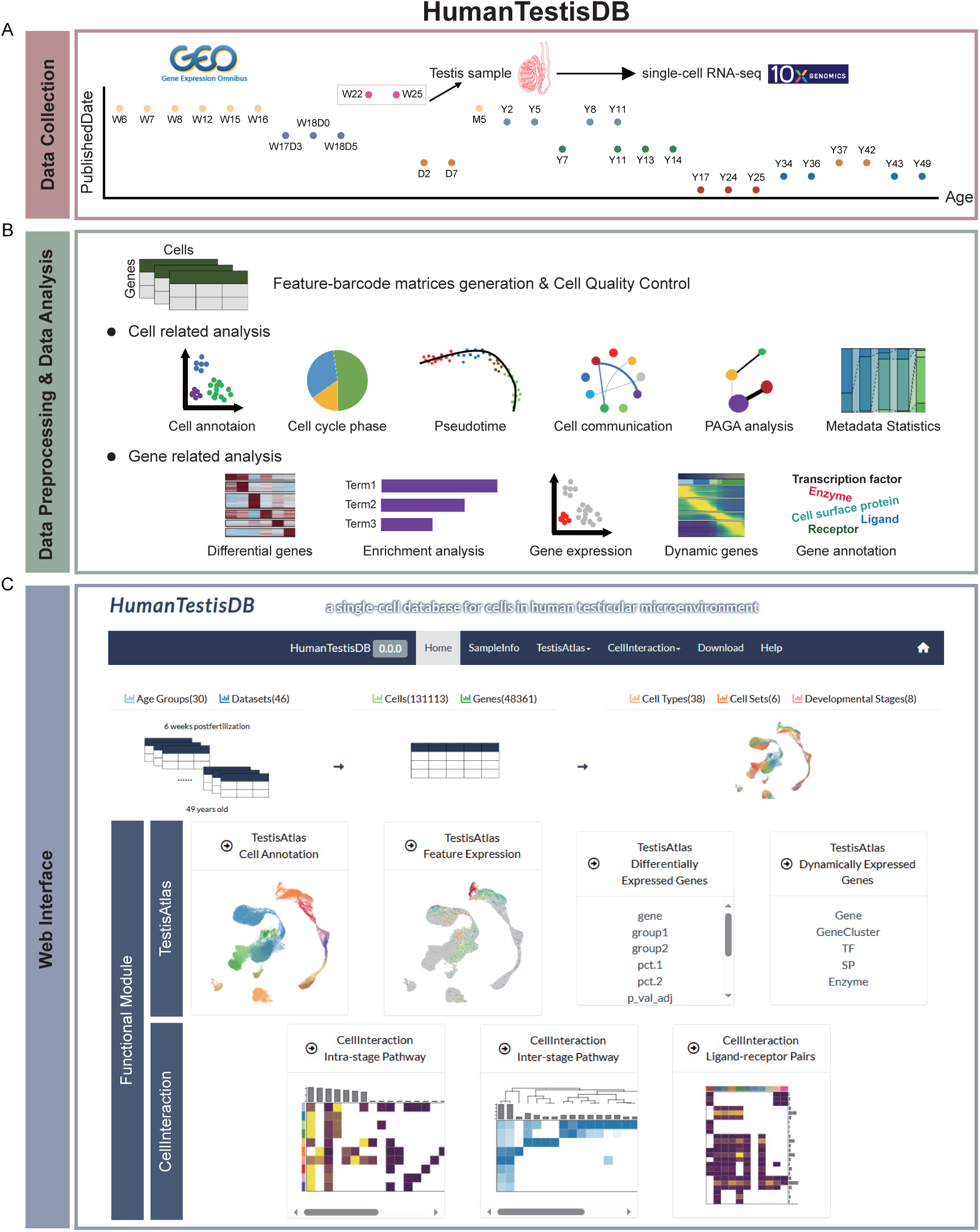
Schematic diagram of the overall design of HumanTestisDB. **A** The sources and sample ages of collected data. W, post-conceptional weeks. D, postnatal days. M, postnatal months. Y, postnatal years. **B** Bioinformatics analysis methods used for processing collected data. **C** The user interface.

**Figure 2.**
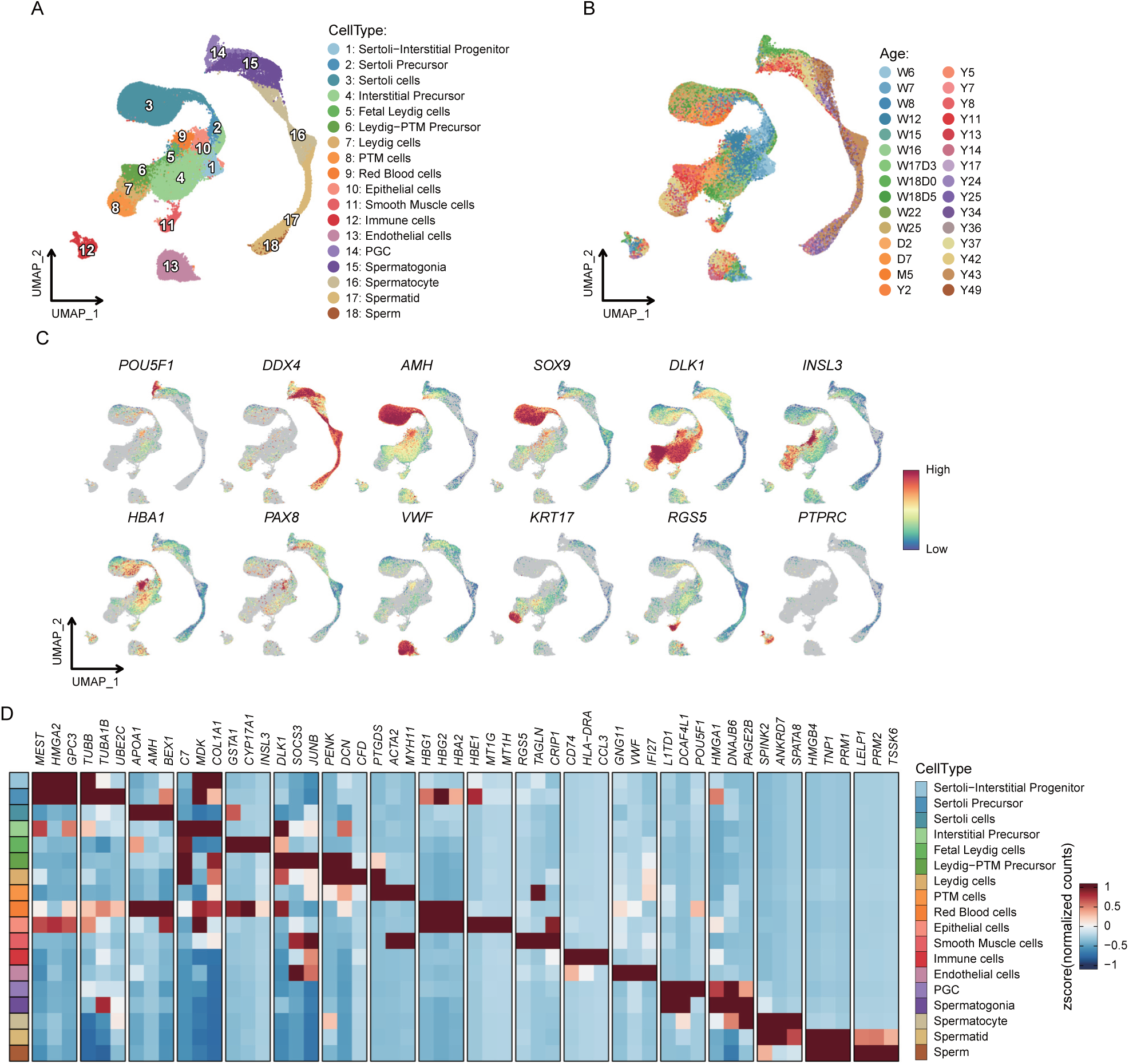
Global transcriptome profiling of all testicular cells in HumanTestisDB. A and. **B** UMAP plot of integrated data from 6 weeks post-fertilization to 49 years old. Each point represents a single cell and is colored according to cell type (A) or age (B). UMAP, uniform manifold approximation and projection. **C** Expression patterns of classical marker genes for major cell types in (A). **D** Heatmap of expression levels for top3 (according to fold change) differentially expressed genes of each cell type in (A). Each row represents a cell type, and each column represents a gene.

Gene Ontology (GO) term enrichment analysis was conducted to confirm our cell type classification, concentrating on the differentially and highly expressed genes in each cell type (Figure S3). The results showed a notable enrichment of GO terms that closely match each cell type’s inherent properties. Specifically, spermatocytes and spermatids have shown significant enrichment in processes associated with meiosis and spermatid differentiation, respectively. Furthermore, Sertoli Precursors demonstrated a substantial correlation with chromosome segregation-related terms, consistent with our finding that most of these cells are in the S/G2M phase of the cell cycle (Figure 4B).

The collection of all 131,113 cells in the database is designated as “AllCells” (Figure 2A). In addition to this large cell set, cells associated with key biological events of spermatogenesis were isolated from the “AllCells” set, forming several subsets named “GermCells”, “GermCells_part1”, “GermCells_part2”, “GermCells_part3”, and “SomaticCells”. “GermCells” comprises all germ cells; “GermCells_part1” includes primordial germ cells (PGCs), spermatogonia, and early spermatocytes; “GermCells_part2” consists of spermatocytes and early spermatids; “GermCells_part3” encompasses spermatids and immature sperm (**Figure 3**A); and “SomaticCells” refers to cells derived from the Sertoli-Interstitial Progenitor lineage (**Figure 4**A). Several analyses were performed on these cell sets (Figure 1B), and the findings were employed to categorize the 46 samples in the database, from W6 to Y49, into eight developmental stages (**Figure 5**A).

**Figure 3.**
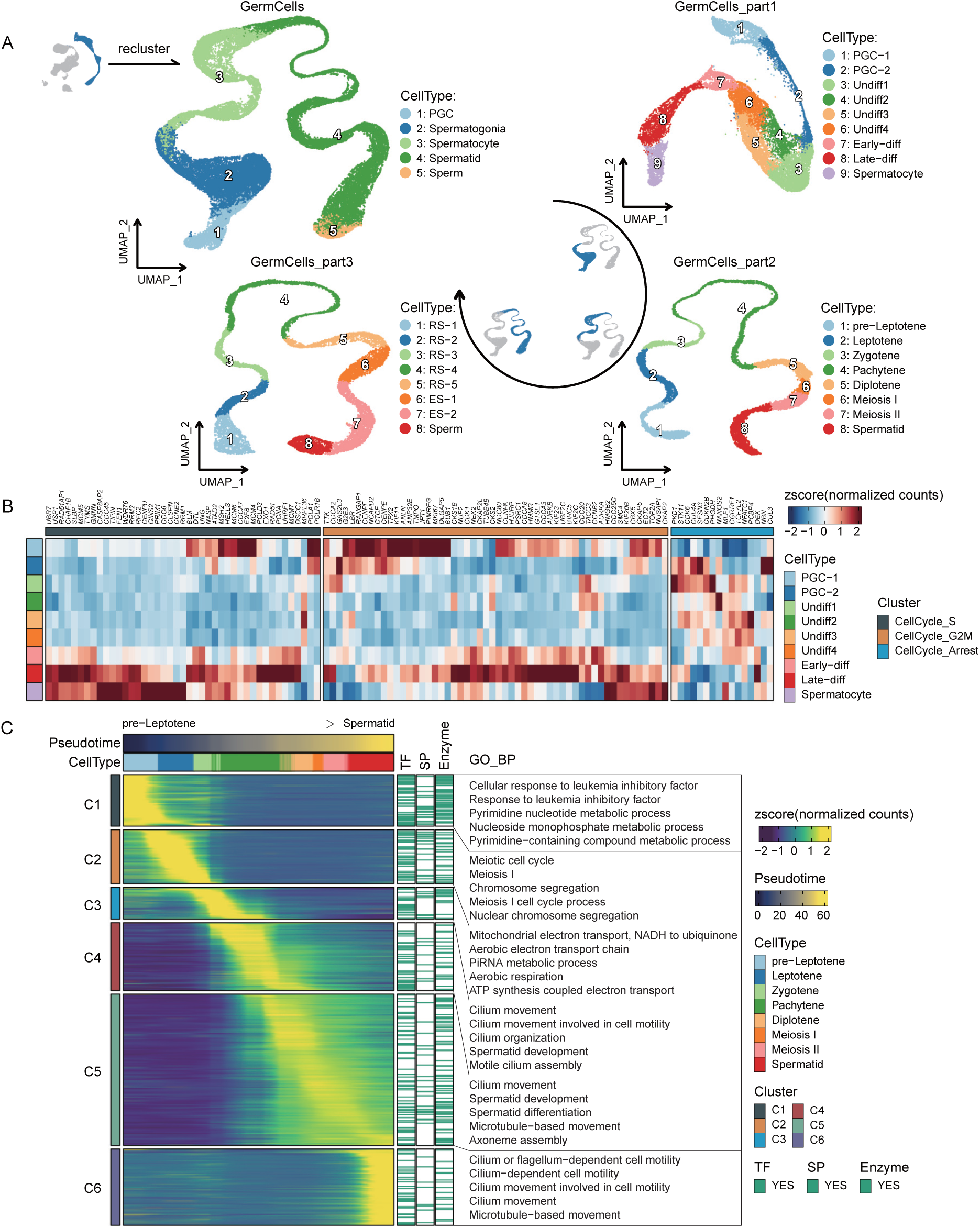
Case study of human testicular germ cells (from PGCs to immature sperm). **A** UMAP plots of germ cells at various spermatogenesis stages, colored by cell type. PGC, primordial germ cell. Undiff, undifferentiated spermatogonia. Early-diff/Late-diff, early/late differentiated spermatogonia. RS, round spermatid. ES, elongated spermatid. **B** The expression levels of cell-cycle-related genes during the transition from PGC-1 to early spermatocytes. **C** Heatmap showing genes dynamically expressed along the developmental trajectory from pre-Leptotene to Spermatid. Genes categorized as transcription factors, cell surface proteins, or enzymes are highlighted in green; associated GO terms are given on the right of the corresponding gene clusters.

We provide our findings via an accessible website comprising six modules: “Home”, “SampleInfo”, “TestisAtlas”, “CellInteraction”, “Download”, and “Help”. The primary functional modules are “TestisAtlas” and “CellInteraction”. The “TestisAtlas” module facilitates comprehensive inquiries into each cell set, encompassing cell attributes, gene expression profiles, differentially expressed genes for each cell type, and genes expressed dynamically throughout cell development. The “CellInteraction” module, in contrast to the “TestisAtlas” module, describes the signaling pathways, ligand-receptor pairs, and cell types involved in the intercellular interactions in the testicular microenvironment at various developmental stages (Figure 1C).

In summary, HumanTestisDB is a database presented through a user-friendly website that contains the results of multiple analyses of the human testis single-cell transcriptome.

### Use case—Proliferation and Differentiation of Germ Cells

HumanTestisDB is primarily utilized to investigate transcriptomic changes throughout spermatogenesis. This database contains four subsets of germ cells: “GermCells”, “GermCells_part1”, “GermCells_part2”, and “GermCells_part3”. “GermCells” delineates the progression from PGC to immature sperm, while “GermCells_part1”, “GermCells_part2”, and “GermCells_part3” outline the three essential stages of spermatogenesis: PGC differentiation and spermatogonia mitosis, spermatocyte meiosis, and spermiogenesis, respectively (Figure 3, Figure S4-6). The “TestisAtlas” module comprises four functional pages: “Cell Annotation”, “Feature Expression”, “Differentially Expressed Genes”, and “Dynamically Expressed Genes” (Figure 1C), allowing users to select the preferred cell set through the “Cell Set” option.

”GermCells_part1” comprises two types of PGCs, six types of spermatogonia, and early spermatocytes (Figure 3A, Figure S4A, B). PGC-1 cells exhibit the expression of pluripotency markers *POU5F1* and *NANOG*, whereas PGC-2 cells lack *POU5F1* expression but express *DDX4* (Figure S4C), aligning with findings documented in prior research about FGCs [15–19]. Analysis of undifferentiated spermatogonia (Undiff) shows that Undiff2 mainly expresses genes related to cytoplasmic translation and ATP synthesis. In contrast, Undiff3 is involved in various biological processes, such as cell cycle progression [20], signal transduction [21, 22], inflammatory responses [23–25], and spermatogonia migration induction [26] (Figure S4E, Table S2). After that, undifferentiated spermatogonia progresses to the differentiation stage and initiates meiosis (Figure S4B). A distinct trend was identified when examining the expression of cell-cycle-related genes in PGC-1 to early spermatocytes (Figure 3B). PGC-1 expressed genes associated with the S and G2M phases, whereas PGC-2 to Undiff4 exhibited increased expression of genes related to cell-cycle arrest compared to PGC-1. In early differentiated spermatogonia to early spermatocytes, the expression of genes associated with cell-cycle arrest diminished. However, the expression of genes pertinent to the S and G2M phases significantly increased (Figure 3B). This aligns with the research indicating that early PGCs are in an active proliferation phase, late PGCs undergo mitotic suppression, and male germ cells do not initiate meiosis until adolescence [4, 27, 28].

HumanTestisDB provides thorough information on spermatocyte stages in “GermCells_part2” that spans pre-Leptotene to Meiosis II, which differs from many other research (Figure 3A). Characteristics of meiosis beginning include the majority of cells in the S phase and increased expression of mitochondrial genes (Figure S5A, C). Furthermore, HumanTestisDB enables users to see significant changes in gene expression patterns throughout these stages, with specific gene clusters indicating participation in various biological processes (Figure 3C). The C1 gene set is mainly expressed in pre-leptotene and participates in the pyrimidine nucleotide metabolic process, which is consistent with the fact that a large amount of DNA is synthesized in pre-leptotene. The C2 gene set is mainly expressed in leptotene and zygotene, and it can be seen that these genes are enriched with terms related to meiosis. Primarily expressed in zygotene and pachytene, the C3 gene set is prominent in ATP synthesis and piRNA metabolism. Kawase M et al. showed that testicular piRNA can be divided into fetal piRNA, pre-pachytene piRNA, and pachytene piRNA [29], indicating the reliability of our enrichment results. The C4 gene set is predominantly expressed during pachytene, a stage characterized by the close association of homologous chromosomes and the exchange and recombination of incomplete DNA segments between alleles. The GO enrichment results demonstrate that the C4 gene set is linked to cilia. Xie H et al. showed that deleting germ cell-specific ciliary genes could increase germ cell apoptosis, reduce crossover formation, and damage double-strand break repair [30]. The C5 gene set is expressed from diplotene to early spermatids, exhibiting the most prominent expression peak across several cell types. These genes are implicated in spermatid development and are enriched in terms associated with sperm structure. The C6 gene set is expressed explicitly in early spermatids, enriching terms associated with sperm structure (Figure 3C).

As observed in “GermCells_part3” (Figure 3A), the last stage, spermiogenesis, entails turning spermatids into immature sperm. According to HumanTestisDB, chromatin condensation and subsequent transcriptional shutdown cause cells’ gene expression to decrease as round spermatids give way to elongated spermatids (Figure S6B), a condition that is also seen in mice [31]. The “TestisAtlas” module’s “Dynamically Expressed Genes” page lists genes that exhibit dynamic expression throughout the transition from round spermatids to immature sperm, grouping them into six clusters based on patterns of expression (Figure S6C). Following their departure from meiosis and subsequent spermatid transformation, early spermatids exhibit differential expression of genes linked to cell division and spermatid development (Figure S5B, Figure S6A, C). Using the gene data from GeneCards, the relationship between each of the 1311 genes in the last five clusters and spermiogenesis-related events was deduced, providing crucial new information for future spermatid biogenesis and function research (Figure S6C, Table S3).

The four germ cell subsets in HumanTestisDB examine the complete process or specific aspects of spermatogenesis, which is crucial for elucidating the molecular mechanisms of sperm production and discovering critical regulatory genes.

### Use case—Key Roles of Sertoli and Leydig Cells in the Seminiferous Tubule Microenvironment and Their Developmental Trajectories

Sertoli and Leydig cells play crucial roles in forming and maintaining the seminiferous tubule microenvironment [32–36]. HumanTestisDB, a critical tool for advancing testicular research, enables the study of germ cells and enhances comprehension of somatic cell differentiation. The “SomaticCells” cell set was created to investigate transcriptomic changes during the differentiation of these somatic cells (Figure 4A). During the critical period of 6 to 7 weeks post-fertilization, the predominant cell type identified was the Sertoli-Interstitial Progenitor (Figure 4C). These progenitors subsequently differentiate into Sertoli or interstitial cells (Figure 4A). Following the commitment to the interstitial cell lineage, an intriguing transition occurs, with cells differentiating into Fetal Leydig, mature Leydig, and PTM cells. The differentiation process is characterized by unique gene expression patterns, which are accessible in the “TestisAtlas” modules (Figure 4D, E, Figure S7).

**Figure 4.**
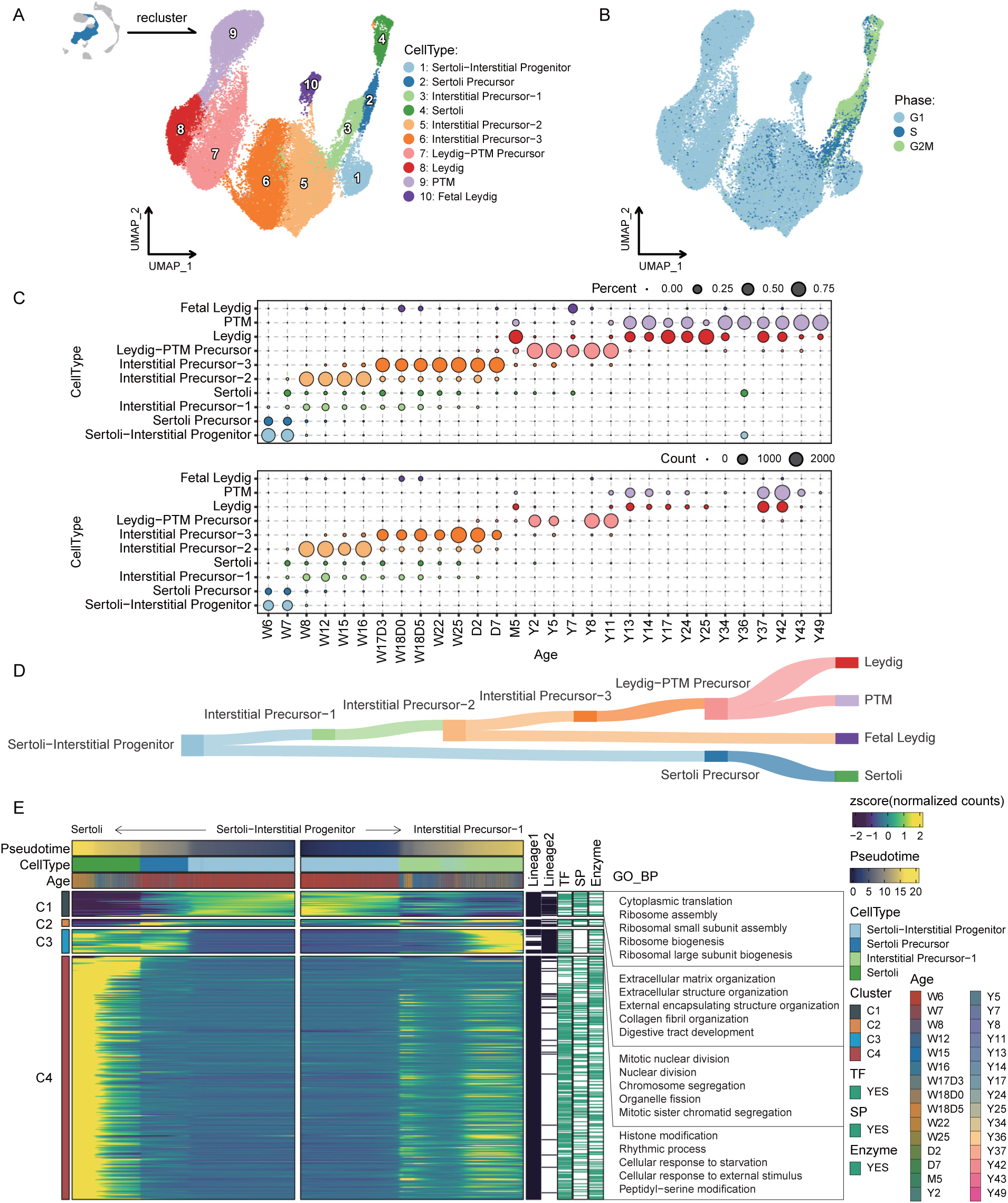
Case study: data analysis focusing on human testicular somatic cells. A and. **B** UMAP plot obtained by re-clustering interstitial-associated cells and early Sertoli cells in Figure 2A, colored by cell type (A) or cell cycle phase (B). **C** Statistical plot showing the distribution of each somatic cell type in each age group. The size of the dots is proportional to the percentage of cells (upper row) or the number of cells (lower row). **D** Schematic illustration of somatic cell development trajectory. **E** Heatmap illustrating dynamically expressed genes along developmental trajectories (from Sertoli-Interstitial Progenitor to Sertoli and from Sertoli-Interstitial Progenitor to Interstitial Precursor-1), with transcription factors, cell surface proteins, or enzymes highlighted in green, and enriched biological processes for each gene cluster displayed.

Investigating the “Cell Annotation” page under the “TestisAtlas” module yielded additional insights (Figure 1C). Among these developing cell types, a distinct age-related distribution pattern was recorded (Figure 4C). In particular, stages W8-W16 and W17D3-D7 are when Interstitial Precursor-2 and Interstitial Precursor-3 are most prevalent. Interestingly, the predominant cell type before puberty is Leydig-PTM Precursor cells. After puberty, these precursors differentiate into Leydig and PTM cells, undergoing a significant shift. Furthermore, most Sertoli Precursors, Interstitial Precursor-1, and early Sertoli cells were discovered to be in the proliferation phase, specifically the S and G2M phases. On the other hand, other cell types mainly inhabit the G1 phase of the cell cycle (Figure 4B).

In conclusion, the “SomaticCells” cell subset offers a thorough illustration of the Sertoli-Interstitial Progenitor’s developmental trajectory, enhancing our comprehension of somatic cell differentiation and providing the groundwork for future research into the traits of somatic cells at various stages.

### Use case—Intricate Cell-Cell Communication in the Testicular Microenvironment Across Developmental Stages

The testis is a complex organ with a variety of cells and a unique physical structure. Since most testicular cell types are functionally interdependent, normal spermatogenesis depends on their interactions [37–39]. HumanTestisDB describes the developmental process from W6 to Y49, during which testicular cell types undergo significant changes, particularly germ and interstitial-derived cells. Based on the distribution of various cell types in these samples, W6-Y49 may be categorized into eight developmental stages: ① W6-W7, the germ cells are all PGC-1, while the interstitial-derived cells are mostly Sertoli-Interstitial Progenitors and a small number are Interstitial Precursors. ② Most of the interstitial-derived cells in stage W8-W16 are Interstitial Precursors, with Interstitial Precursor-2 being the predominant one, whereas a tiny number of PGC-2 is visible compared to stage W6-W7. ③ W17D3-W25, when compared to W8-W16, there are many more PGC-2 cells and a few spermatogonia. Most interstitial-derived cells are Interstitial Precursors, mostly Interstitial Precursor-3. ④ D2-D7, compared with stage W17D3-W25, there are significant changes in the distribution of endothelial and smooth muscle cells. ⑤ In stage Y2-Y8, in contrast to stage D2-D7, PGCs are absent, most germ cells are undifferentiated spermatogonia, with a limited presence of differentiated spermatogonia. On the other hand, most interstitial-derived cells are Leydig-PTM precursors. ⑥ In stage Y11, there is an increase in the number of differentiated spermatogonia compared to stage Y2-Y8, along with the emergence of a limited number of spermatocytes. Simultaneously, most interstitial-derived cells are Leydig-PTM precursors, with a limited presence of Leydig and PTM cells. ⑦ In Y13, relative to stage Y11, a limited quantity of spermatids is present, whereas the interstitial-derived cells consist of Leydig and PTM cells. ⑧ In stage Y14-Y49, in contrast to stage Y13, there exists a diverse array of germ cells, encompassing spermatogonia to immature sperm, while interstitial-derived cells include Leydig and PTM cells (Figure 5A, Figure S8, Figure S9).

**Figure 5.**
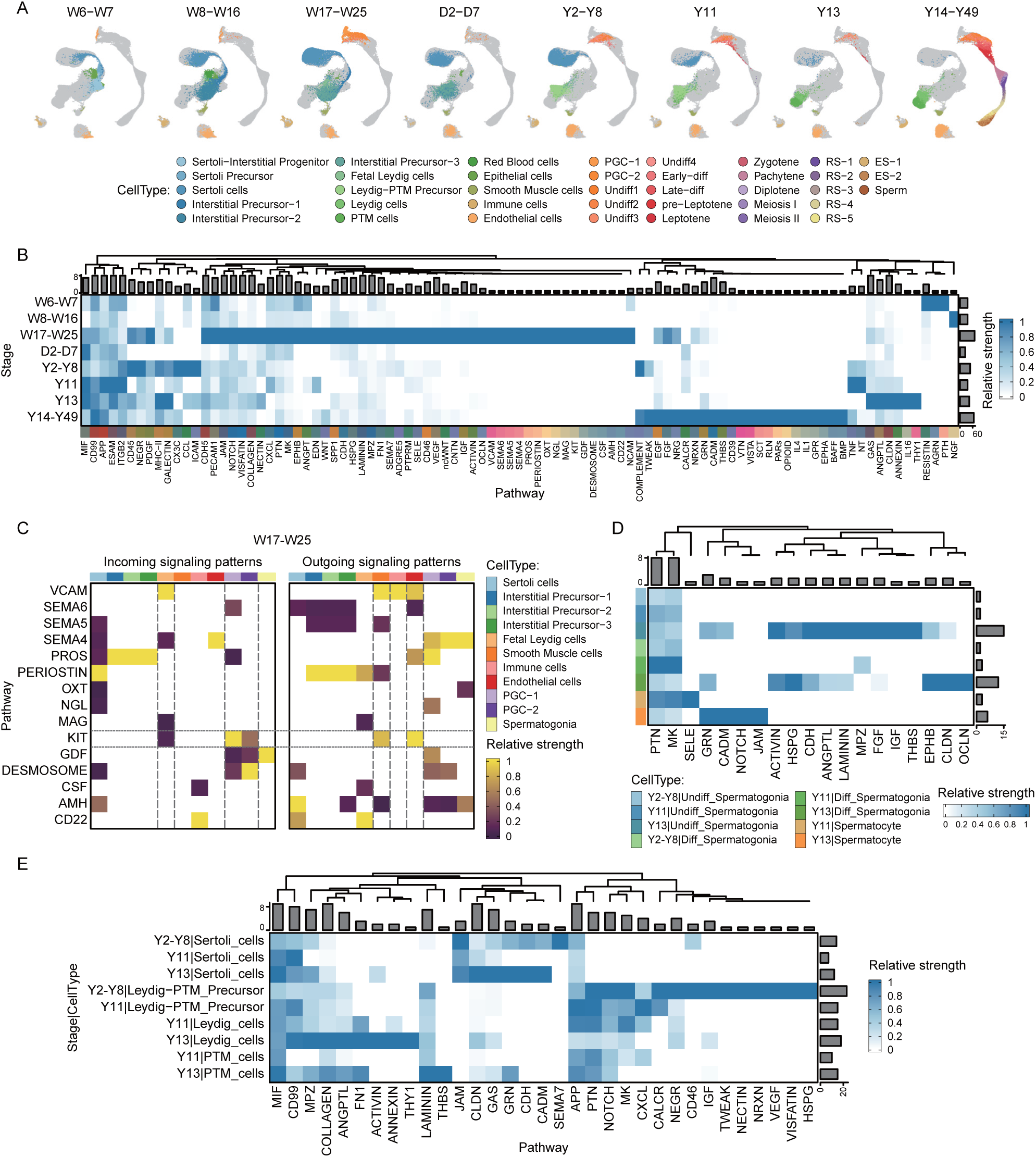
Cell–cell communication dynamics within the testicular microenvironment. **A** UMAP plots of eight distinct developmental stages that differ in cell type composition, colored by cell type. **B** Heatmap depicting relative strength of signaling pathways common or specific to stages, with normalized columns for comparability. Each row represents a stage, and each column represents a pathway. The height of the bars at the top represents the number of stages in which each pathway is detected. The width of the bars on the right represents the number of pathways detected within each stage. **C** Heatmap illustrating receivers and senders mediating cell-cell communications specific to stage W17-W25, with normalized columns for signal strength comparability. Each row represents a pathway, and each column represents a cell type. **D** Heatmap presenting communication strength of signaling pathway received by different germ cell types at specified stages, with normalized columns for comparability. Each row represents a cell type, and each column represents a pathway. The height of the bars at the top represents the number of cell types that receive each pathway. The width of the bars on the right represents the number of pathways received by each cell type. **E** Heatmap showcasing communication strength of signaling pathway sent by different somatic cell types at specified stages, with normalized columns for comparability. Each row represents a cell type, and each column represents a pathway. The height of the bars at the top represents the number of cell types that receive each pathway. The width of the bars on the right represents the number of pathways received by each cell type.

This study investigated cell-cell interactions at each developmental stage utilizing the CellChat software. Specific signaling pathways, including CD99, APP, and ESCM, demonstrate sustained activity across all eight stages (Figure 5B). It also uncovers stage-specific pathways, for example, 15 different pathways, including VCAM and CD22, are primarily active during W17-W25 (Figure 5B).

The “Intra-stage Pathway” page comprehensively outlines the signaling pathways identified at each developmental stage and the associated sender and receiver cells. Compared to “Intra-stage Pathway”, “Ligand-receptor Pairs” explores more deeply, providing detailed insights into the ligand-receptor pairs (Figure 1C). Concentrating on stage W17-W25, HumanTestisDB validates prior studies, emphasizing that PGCs express the *KIT* gene, a transmembrane tyrosine kinase receptor [40–45]. Furthermore, it demonstrates that *KITL*, the ligand for *KIT*, is mainly expressed by endothelial and smooth muscle cells, and Fetal Leydig cells are the principal recipients of KIT signaling (Figure 5C). This observation confirms the essential function of *KITL*-*KIT* signaling in developing and maintaining Fetal Leydig cells [46].

Unlike the “Intra-stage Pathway” and “Ligand-receptor Pairs” pages, the “Inter-stage Pathway” page allows for cross-stage comparisons of signals received or sent by cell types (Figure 1C). For example, during meiosis, a process that begins at the commencement of puberty in males and is critical for producing the large number of gametes required for male fertility, HumanTestisDB shows a different development of germ cells into spermatocytes at stages Y11 and Y13, as opposed to stage Y2-Y8. Specifically, stage Y11, unlike stage Y13, continues to include Leydig-PTM precursors (Figure 5A, Figure S8, Figure S9). The present study compared the signals received by germ cells and those sent by the primary somatic cells during these three stages (Figure 5D, E). Analysis of signal strength suggests that spermatogonia at stage Y13 receive a more comprehensive range of signals than their earlier-stage counterparts, including ACTIVIN, IGF, and EPHB (Figure 5D). Notably, research reveals that *IGF1* plays an essential function in controlling spermatogenesis and encouraging the differentiation of spermatogonia into primary spermatocytes [47–49].

In conclusion, HumanTestisDB provides a verifiable repository of signaling pathways, thereby facilitating advanced in vitro germ cell culture research.

## Discussion

HumanTestisDB is a comprehensive database, however, its scope is constrained by the limited variety of datasets available. A main limitation is the underrepresentation of Sertoli cells in samples from puberty to adulthood. According to Guo J et al., mature Sertoli cells generally exceed the size range effectively recorded by 10x Genomics technology, leading to an inadequately low representation of adult Sertoli cells in the database [5]. This highlights the necessity of incorporating more comprehensive Sertoli cell sequencing data, which would significantly enhance the content and functionality of HumanTestisDB.

Future database upgrades should incorporate a broader range of demographic data to increase relevance and applicability. As single-cell technologies progress, it is essential to keep the database updated by including new findings and methodological innovations. These upgrades would enhance the database’s research applicability and enable the translation of these findings into clinical settings, especially regarding male reproductive health and the management of associated disorders.

Furthermore, studying the epigenetic variables influencing testicular development is a promising but unexplored field. Exploring this area of testicular transcriptomics is a promising avenue for future study, with the potential to identify new treatment targets and mechanisms in male reproductive health.

## Conclusion

An important step forward in understanding the complexities of the human testicular microenvironment is represented by HumanTestisDB. The fact that it was created highlights the growing need for integrative databases to summarize and interpret biological systems’ complex nature. By combining disparate data, HumanTestisDB goes beyond being a simple archive and improves our understanding of testicular biology. It provides a comprehensive transcriptome panorama of the human testis, following its development from embryonic to adulthood, by integrating 46 single-cell sequencing datasets.

The careful identification of 38 different cell types and the explanation of their developmental trajectories form the basis of HumanTestisDB’s originality. This discovery is revolutionary because it provides previously unheard-of insights into the cellular mosaic of human tissues. Furthermore, the database plays a crucial role in deciphering the complex communication network that permeates different testicular cells at various stages of development. The focus on developmental stratification is particularly significant, reflecting the changing viewpoint in developmental biology that recognizes cell states’ fluid and dynamic characteristics. Its contributions are essential in promoting further research, potentially revealing novel reproductive health and disease treatment avenues.

## Materials and methods

### Data collection and preprocessing

Among the 46 datasets included in HumanTestisDB, 3 datasets (W22, W25A, and W25B) were generated by our laboratory through performing 10x Genomics sequencing on donated testicular samples, while the other 43 datasets were sourced from published studies. The literature-based data was identified using keywords like “human testis”, “scRNAseq”, and “10x Genomics”. From these studies, we obtained SRA files, converted them to FASTQ format using the fastq-dump command from sratoolkit-2.10.8 [50], and processed them with Cell Ranger 5.0.0 [51] to generate feature-barcode matrices. When FASTQ files were unavailable, we directly acquired feature-barcode matrices from the publications. The laboratory-generated data were similarly processed with Cell Ranger 5.0.0.

### Human fetal gonads preparation for single cell RNA sequencing

Male fetal gonads were washed three times with PBS, divided into smaller sizes (around 2 mm each) using scissors. Single testicular cells were obtained using two-step enzymatic digestion. Briefly, tissues were treated with collagenase type IV (GIBCO cat # 17104) for 10-20 min and 0.25% EDTA (GIBCO cat # 25200056) for 3-8 min at 37°C. Then, the digestion was then stopped by adding 10% FBS (GIBCO cat # 10082147). Single testicular cells were obtained by filtering through 70 μm (Fisher Scientific cat # 08-771-2) and 40 μm (Fisher Scientific cat # 08-771-1) strainers. The cells were pelleted by centrifugation at 600-800 g for 5 min, and wash with PBS twice. Cell number was counted using cell counter (C100-SE), and the cells were then resuspended in PBS + 0.4% BSA (Thermo Fisher Scientific cat # AM2616) at a concentration of ∼1,000 cells/uL ready for single-cell sequencing.

### Single cell RNA-seq performance, library preparation and sequencing

We aimed to capture ∼4,000-5,000 cells. Briefly, cells were diluted following manufacturer’s instructions, and 70 uL of total mixed buffer together with cells were loaded into 10x Chromium Controller using the Chromium Single Cell 3′ v3.1 reagents (10X genomics, PN-1000121). In brief, single-cell suspensions were used for GEMs generation, GEM-RT, cDNA amplification and purification. cDNA concentrations were measured with a Qubit dsDNA HS and BR Assay Kits (Thermo Fisher Scientific, Q32584). Indexed libraries were constructed according to the user guide. Post library construction QC were analysed on a Fragment Analyzer or LabChip, and the library product were sequenced on a illumina NovaSeq 6000 sequencer at the Nanjing Jiangbei New Area Biopharmaceutical public service platform (Nanjing, China) with the following sequencing strategy: 150-bp read length for paired-end.

### Data quality control

Each feature-barcode matrix was converted into a Seurat object using the CreateSeuratObject function from the SeuratObject [52] package. Quality control was conducted using the RunCellQC function of the SCP [53] package. We applied scDblFinder [54] for doublet detection, for the reason that it is common to have 10-20% doublets in single-cell experiments. Low-quality cells were filtered out based on median absolute deviation (MAD) [55]. The retained cells met the following standards: percent.mt < 20, percent.ribo < 50, nFeature_RNA > 500, and nCount_RNA > 1000.

### Batch effect correction

To mitigate batch effects from different sources, we first reproduced the dimensionality reduction results of each literature to determine the magnitude of batch effects between the datasets. We divided 46 datasets into 14 groups, namely: Group1 (W6, W7, W8, W12, W15, W16), Group2 (W17D3, W18D0, W18D5), Group3 (W22, W25A, W25B), Group4 (D2-I, D2-T, D7-I, D7-T), Group5 (M5), Group6 (Y7-rep1, Y7-rep2), Group 7 (Y2, Y5, Y8), Group8 (Y17-1-rep1, Y17-1-rep2, Y24-rep1, Y24-rep2, Y25-rep1, Y25-rep2), Group9 (Y11-1-rep1, Y11-1-rep2, Y13-rep1, Y13-rep2, Y14-rep1, Y14-rep2), Group10 (Y11-2), Group11 (Y43-2-Spc, Y43-2-Spd), Group12 (Y34-1, Y36-1, Y49-1), Group13 (Y43-1-Spc, Y43-1-Spd), and Group14 (Y37-2-I, Y37-2-T, Y42-1-I, Y42-1-T). We pass the grouping information to the “batch” parameter of the Integration_SCP function in the SCP package and set nHVF = 870 and integration_method = “CSS” [56] for data integration.

### Cell type identification

Cell types were deduced from the integrated data based on a combination of criteria: classical cell type-specific markers (e.g., *AMH* for Sertoli cells, *DDX4* for germ cells), annotations from publications (e.g., Figure S4D), and additional factors like cell cycle phase and gene content. Gene Ontology [57] term enrichment analys is also aided in this identification process.

### Differentially expressed genes calculation

Differentially expressed genes of each cell type were identified using the RunDEtest function of the SCP packages with default parameters (test.use = “wilcox”, fc.threshold = 1.5, base = 2, min.pct = 0.1, p.adjust.method = “bonferroni”), taking all genes with p_val_adj < 0.05 as differentially expressed genes.

### Dynamically expressed genes calculation

Firstly, the RunSlingshot function of the SCP package was used to compute the pseudotime of cells. The parameters “reduction”, “group.by”, and “start” were used to specify the nonlinear dimensionality reduction graph, cell group, and the cell group to serve as the starting point, respectively. Secondly, pass the calculated pseudotime to the “lineages” parameter of the RunDynamicFeatures function in the SCP package, and set the value of n_candidates to 5000 to calculate the dynamically expressed genes.

### Gene set enrichment analysis and gene annotation

The RunEnrichment function of the SCP package was used for gene set enrichment analysis, using GO terms and the Reactome pathways as background gene sets (db = c(”GO_BP”, “Reactome”), minGSSize = 10, maxGSSize = 500, GO_simplify = TRUE, GO_simplify_padjustCutoff = 0.2, simplify_method = “Rel”, simplify_similarityCutoff = 0.7). The transcription factors, cell surface proteins, and enzymes used for gene annotation were obtained through the PrepareDB function of the SCP package (species = “Homo_sapiens”, db = “TF” or “SP” or “Enzyme”) and are available on the “Download” module of HumanTestisDB.

### Cell cycle related analysis and cell relationships inference

Cell cycle phases were determined using the CellCycleScoring function of Seurat [58] package. The genes related to S/G2M phase were obtained using the CC_GenePrefetch function of SCP packages. The cell-cycle-arrest-related genes were obtained from Li L et al. [4]. These cell-cycle-related genes are also available on the “Download” module of HumanTestisDB. The RunPAGA function of SCP package was used to infer the strength of the correlation between cells, with parameters “group_by”, “linear_reduction”, and “ nonlinear_reduction” used to specify cell group, linear dimensionality reduction graph, and nonlinear dimensionality reduction graph, respectively.

### Cell communication

Before analyzing cell-cell interactions, we refined cell type naming for accuracy. Criteria for retaining cell types included a proportion greater than 1.3% for continuously differentiating cells and a cell count above 10 for terminally differentiated cells. Red blood cells and epithelial cells were excluded. For each developmental stage, cell-cell communication was then analyzed using the CellChat [59] package, as instructed in the tutorial (https://htmlpreview.github.io/?https://github.com/jinworks/CellChat/blob/master/tutorial/CellChat-vignette.html), with all identified communications retained. When comparing cell communication across stages, we extracted the results of each stage and merged them into a matrix. Based on the analysis purpose, we divide each value in the matrix by the row or column sums to obtain the relative communication strength. In addition, the term “Incoming signaling patterns” and “Outgoing signaling patterns” involved in pathway analysis are defined by the CellChat package, as stated by Jin S et al. [60], “Outgoing patterns reveal how sender cells (that is, cells acting as signal sources) coordinate with each other, as well as how they coordinate with certain signaling pathways to drive communication. Incoming patterns show how target cells (that is, cells acting as signal receivers) coordinate with each other to respond to incoming signals.”.

### Database construction and implementation

HumanTestisDB was developed in a container based on docker’s r-base image [61]. The container’s environment was configured using apt and pip. In this loosely isolated environment, website was constructed with Django [62], in which data analysis was performed using R programming language, while backend to frontend interaction was managed using Python and HTML. Accessible at https://shalab.njmu.edu.cn/humantestisdb, the database provides a user-friendly interface for exploring cell type distribution, gene expression patterns, and cell-cell interactions.

### Ethical statement

Written consent forms were received from all donors. The fetal gonad specimens were collected from terminated pregnancies, in compliance with the Nanjing Maternity and Child Health Care Hospital’s Ethical Committee’s approval (Approval No. (2017)68). Additionally, the use of human samples in our experiments received the endorsement of the Ethical Committee of Nanjing Medical University (Approval No. (2017)582). Exclusion criteria for the fetal gonad samples included any signs of deterioration or congestion, as well as the presence of genetic abnormalities or infectious diseases, such as Hepatitis B, AIDS, and syphilis.

## Data availability

The scRNAseq data of testis obtained from donors with gestational ages of 22, 25 weeks are newly-generated and have been deposited in the Genome Sequence Archive, China National Center for Bioinformation / Beijing Institute of Genomics, Chinese Academy of Sciences (GSA: HRA008838) that are publicly accessible at https://ngdc.cncb.ac.cn/gsa [63, 64]. The accession number for other sequencing data is shown in Table S1. The HumanTestisDB database is freely available for non-commercial use at https://shalab.njmu.edu.cn/humantestisdb. Users are able to access any data and visualizations without registration or login.

## CRediT author statement

**Mengjie Wang**: Investigation, Data curation, Formal analysis, Methodology, Visualization, Writing - original draft. **Laihua Li**: Resources, Writing - original draft. **Qing Cheng**: Resources. **Hao Zhang**: Conceptualization, Methodology. **Zhaode Liu**: Resources. **Yiqiang Cui**: Resources. **Jiahao Sha**: Conceptualization, Funding acquisition, Project administration, Resources, Supervision, Writing - review & editing. **Yan Yuan**: Conceptualization, Funding acquisition, Project administration, Supervision, Writing - review & editing. All authors read and approved the final manuscript.

## Acknowledgements

This work was supported by National Natural Science Foundation of China (grant 82122025 (Y.Y.), 82221005 (J.S.)) and National Key R&D Program (grant 2022YFC2702800 (Y.Y.), 2021YFC2700302 (Y.Y.), 2021YFC2700200 (Y.C.)). We acknowledge the authors from published studies to share their single-cell RNA-seq data on human testis samples.

## Figure legends

**Figure S1.**
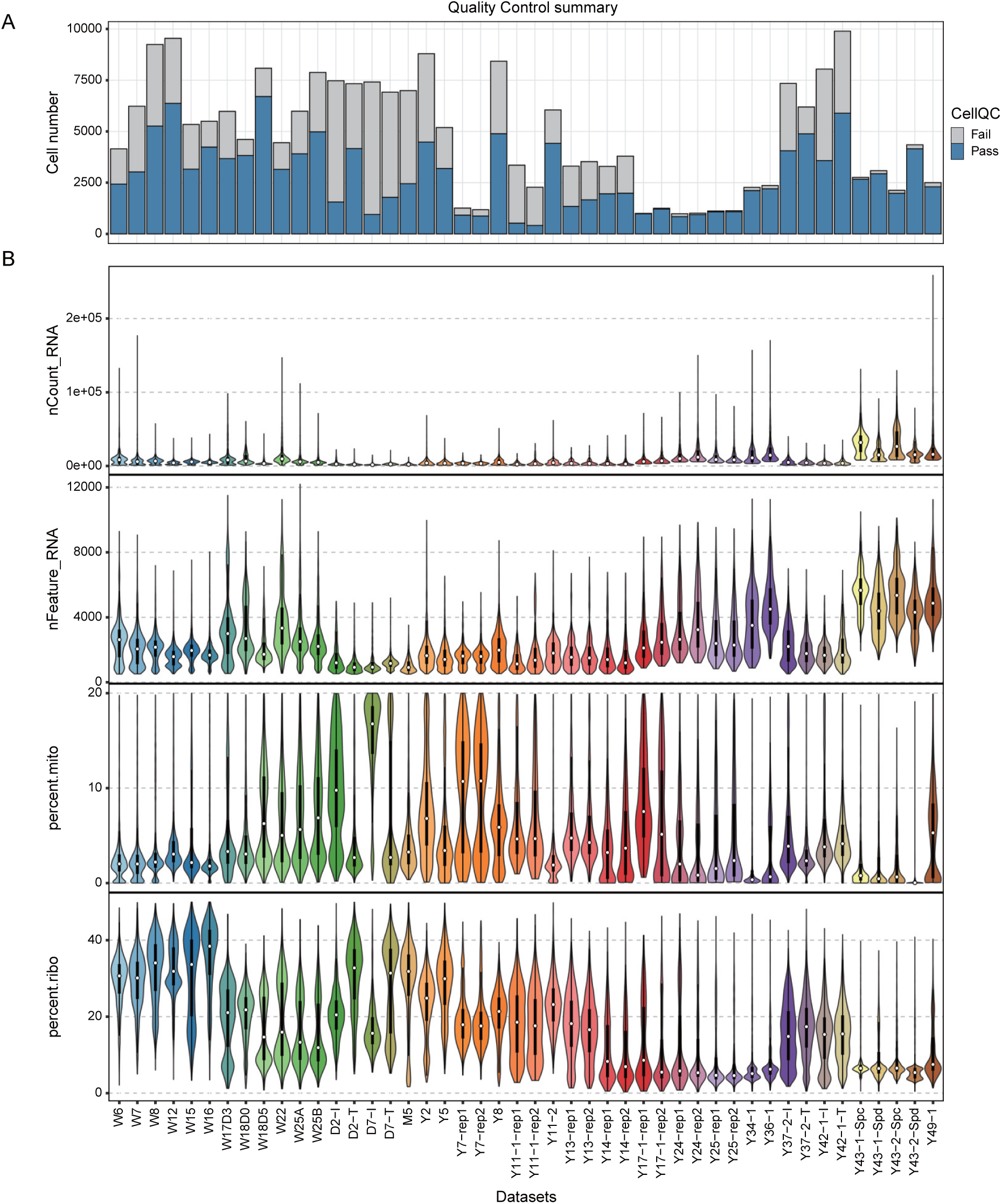
Quality control of single-cell RNA-seq datasets. **A** Bar plot displaying the number of cells passing and failing quality control for each dataset. **B** Violin plots depicting single-cell RNA-seq quality metrics: total molecules detected per cell (nCount_RNA), gene count per cell (nFeature_RNA), mitochondrial gene percentage (percent.mito), and ribosomal gene percentage (percent.ribo).

**Figure S2.**
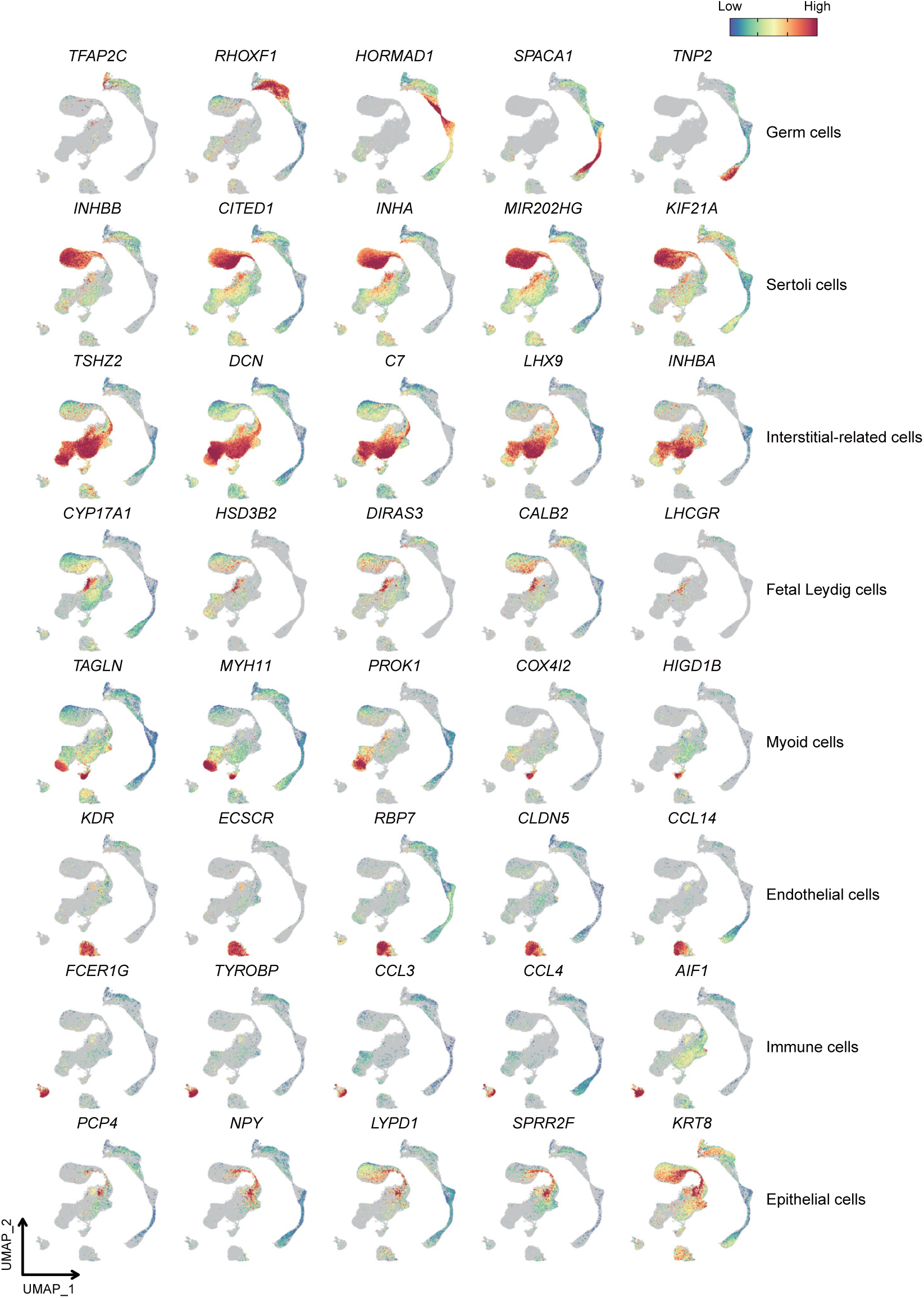
Expression patterns of additional representative markers related to cell types in Figure 2A.

**Figure S3.**
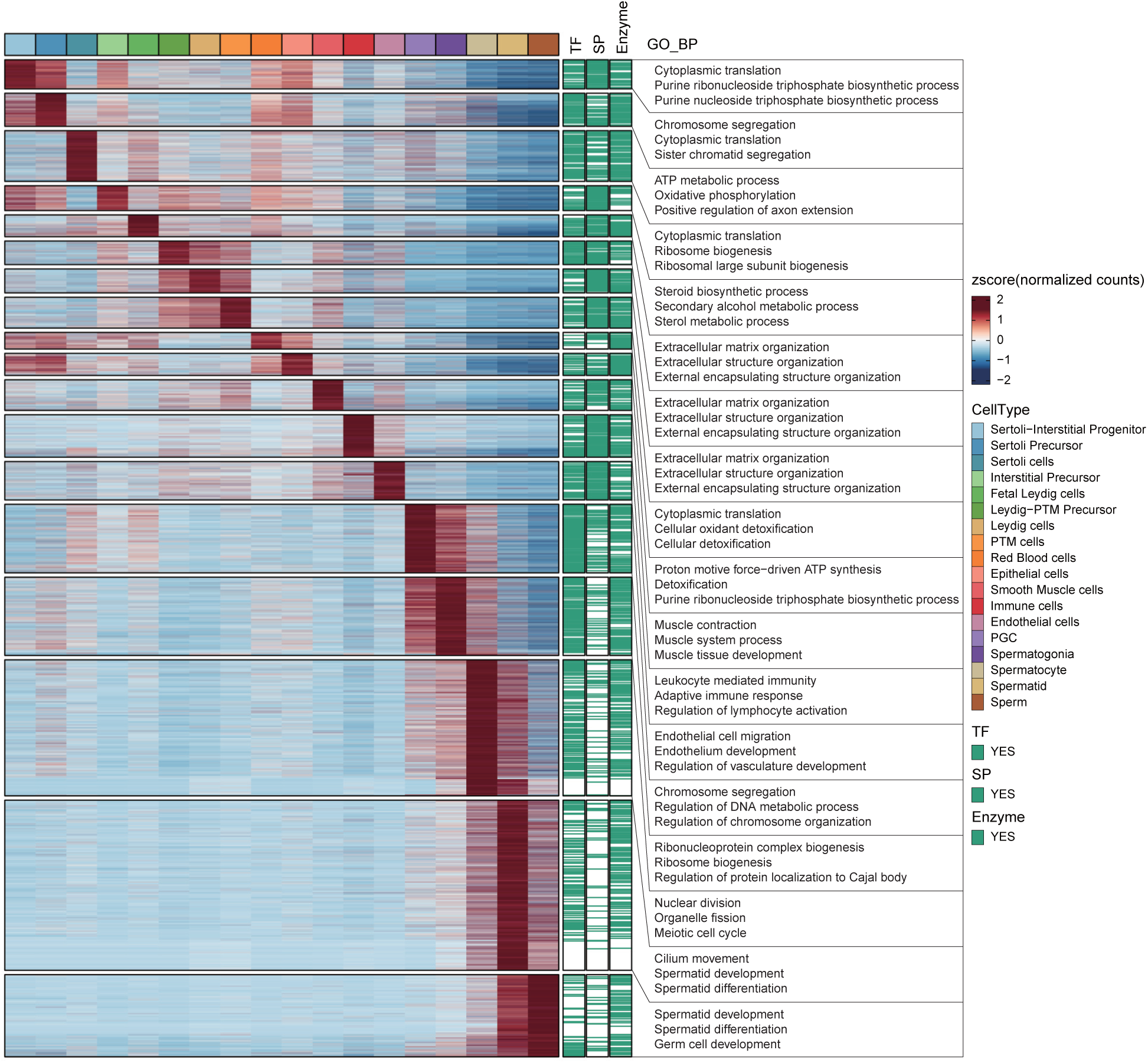
Gene expression difference across cell types in Figure 2A. Heatmap showing differentially expressed genes, with regions corresponding to transcription factors, cell surface proteins, or enzymes marked in green, and enriched biological processes for each gene cluster displayed.

**Figure S4.**
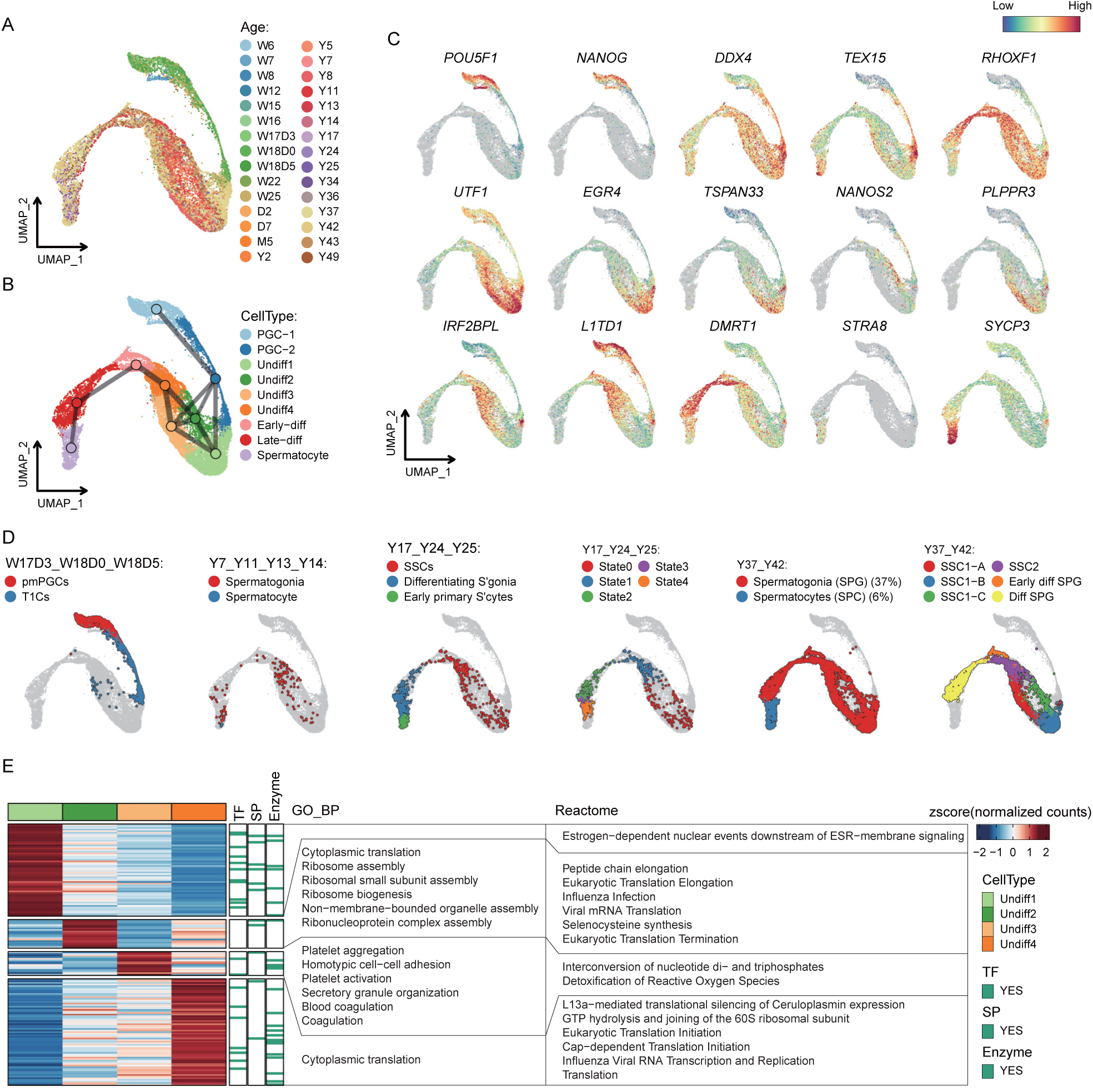
Characterization of the process from PGC to early Spermatocyte. **A** and **B** UMAP plot of the “GermCells_part1” cell set in Figure 3A, colored by age (A) or cell type (B). Line thickness between cell types in (B) indicates connection strength. **C** Expression patterns of classical markers for cell types in (B). **D** Cell annotation from different literature projected on the UMAP plot of the “GermCells_part1” cell set. **E** Heatmap showing differentially expressed genes of undifferentiated spermatogonia in (B), with transcription factors, cell surface proteins, or enzymes highlighted in green, and enriched GO biological processes and Reactome pathways for each gene cluster displayed.

**Figure S5.**
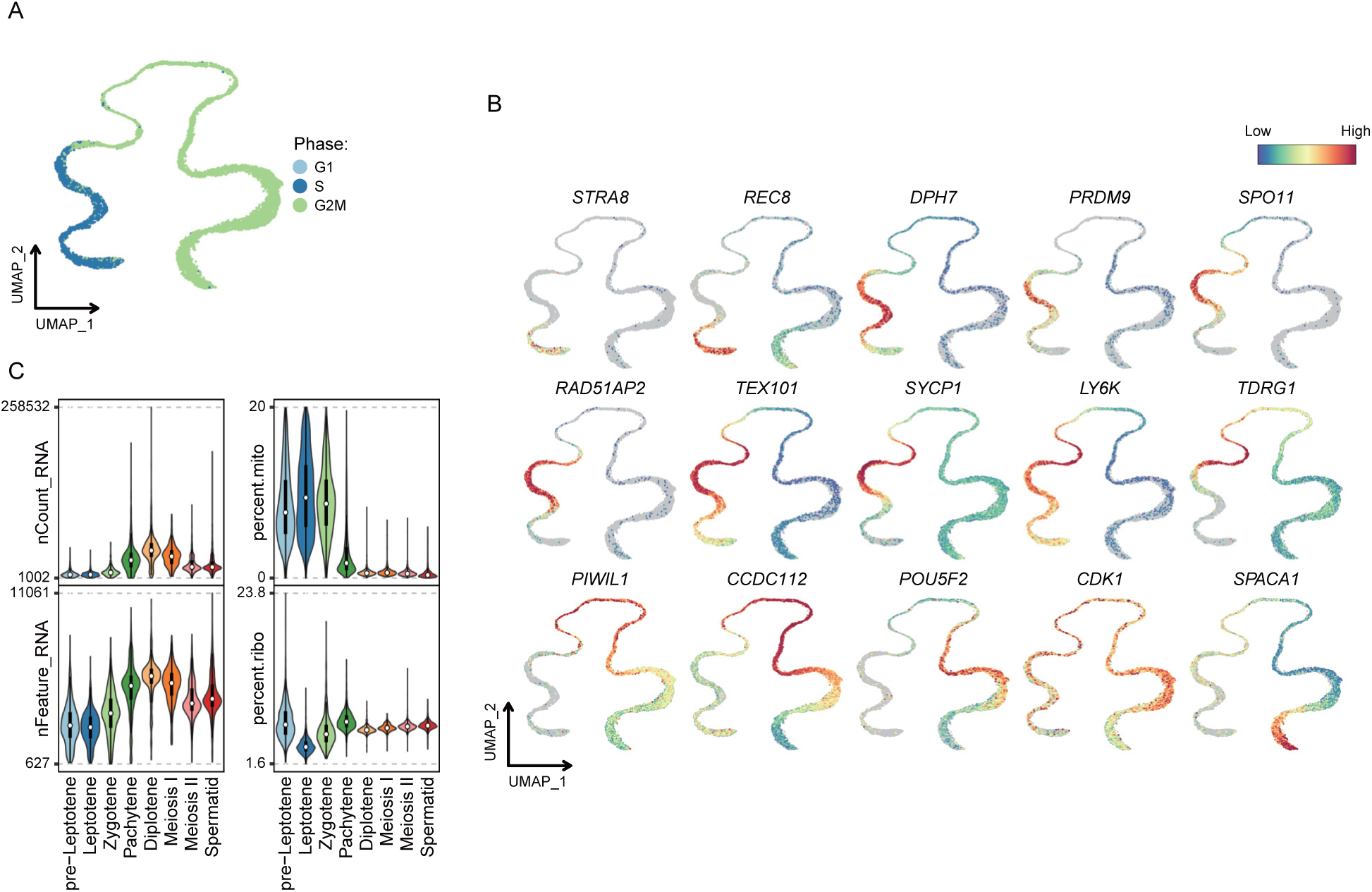
Dynamics during the differentiation from pre-Leptotene to early Spermatid. **A** UMAP plot of the “GermCells_part2” cell set in Figure 3A, colored according to cell cycle phase. **B** Expression patterns of classical markers for cell types in the “GermCells_part2” cell set. **C** Violin plots showing nCount_RNA, nFeature_RNA, percent.mito, and percent.ribo of each cell type in the “GermCells_part2” cell set.

**Figure S6.**
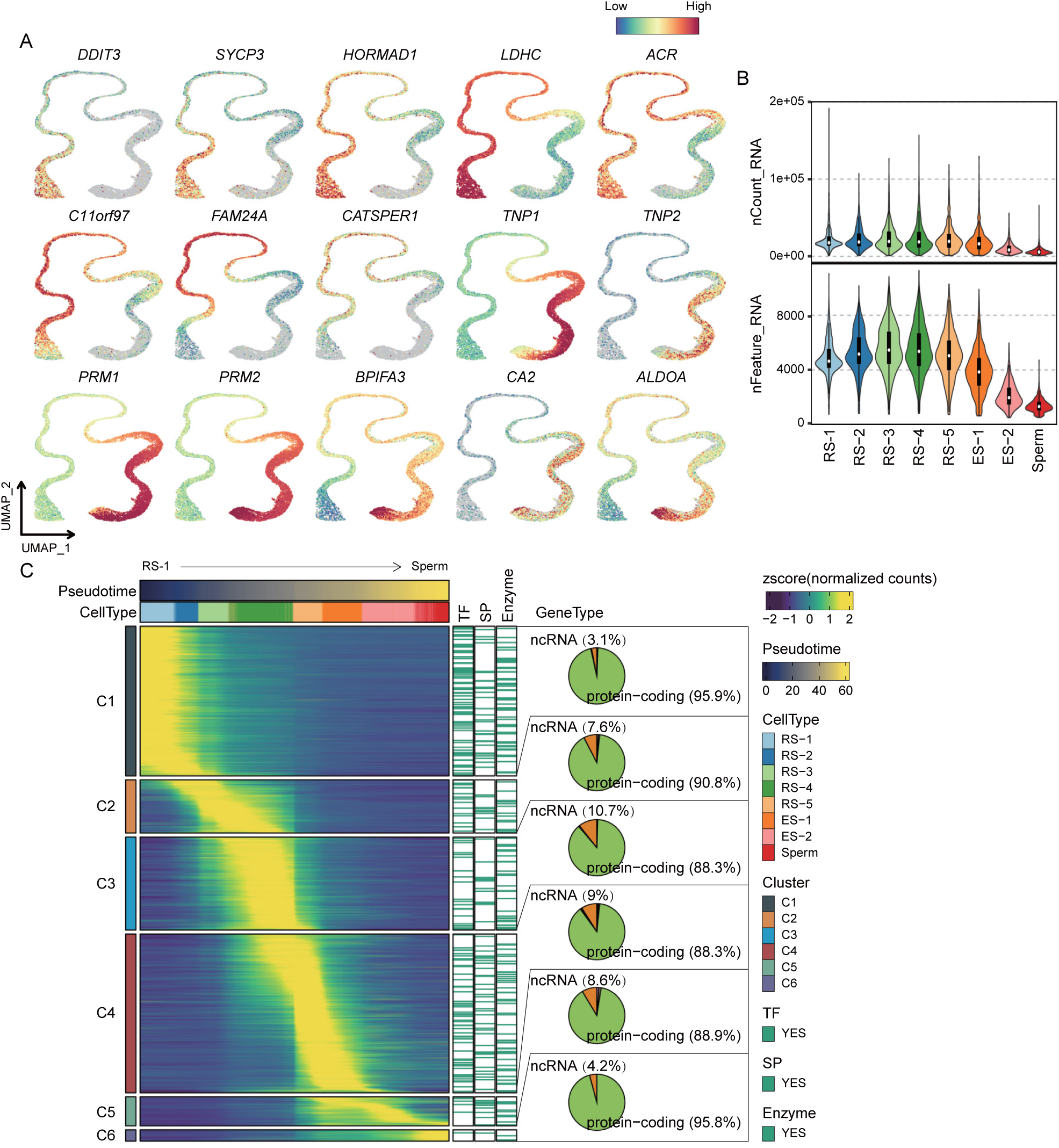
Gene expression dynamics throughout spermiogenesis. **A** Expression profiles of classical markers for cell types in the “GermCells_part3” cell set. **B** Violin plots showing nCount_RNA and nFeature_RNA of each cell type in the “GermCells_part3” cell set. **C** Heatmap illustrating genes dynamically expressed along the developmental trajectory from RS-1 to Sperm, with regions corresponding to transcription factors, cell surface proteins, or enzymes marked in green. The gene types included in each gene cluster are also displayed.

**Figure S7.**
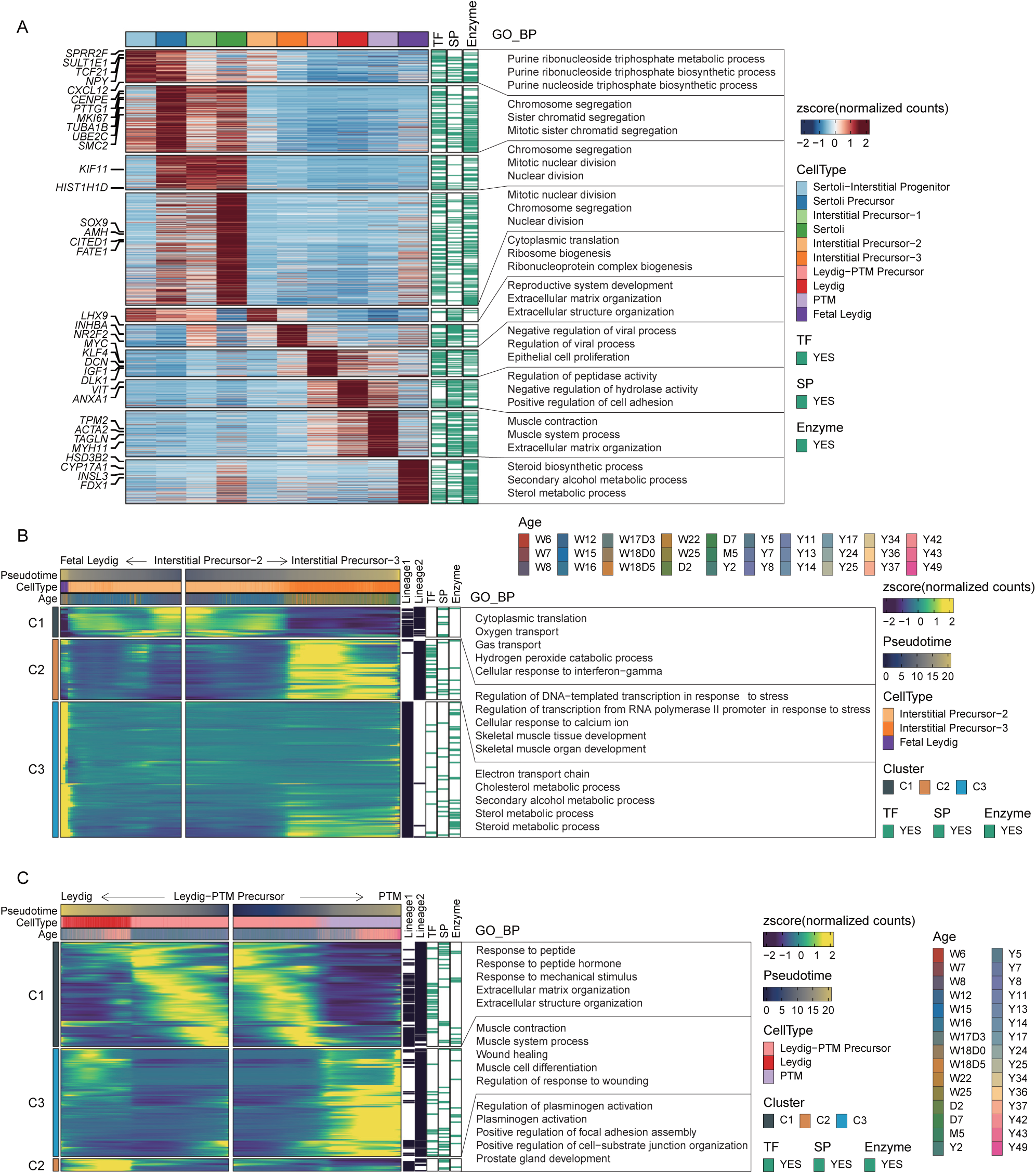
Gene Expression insights and dynamics in somatic cells. **A** Heatmap showing differentially expressed genes for each cell type in Figure 4A, with transcription factors, cell surface proteins, or enzymes highlighted in green. Enriched biological processes for each gene cluster are also displayed. **B and C** Heatmap illustrating genes dynamically expressed along developmental trajectories, with transcription factors, cell surface proteins, or enzymes highlighted in green. Enriched biological processes for each gene cluster are also displayed. From Interstitial Precursor-2 to Interstitial Precursor-3 and from Interstitial Precursor-2 to Fetal Leydig (B). From Leydig-PTM Precursor to Leydig and from Leydig-PTM Precursor to PTM (C).

**Figure S8.**
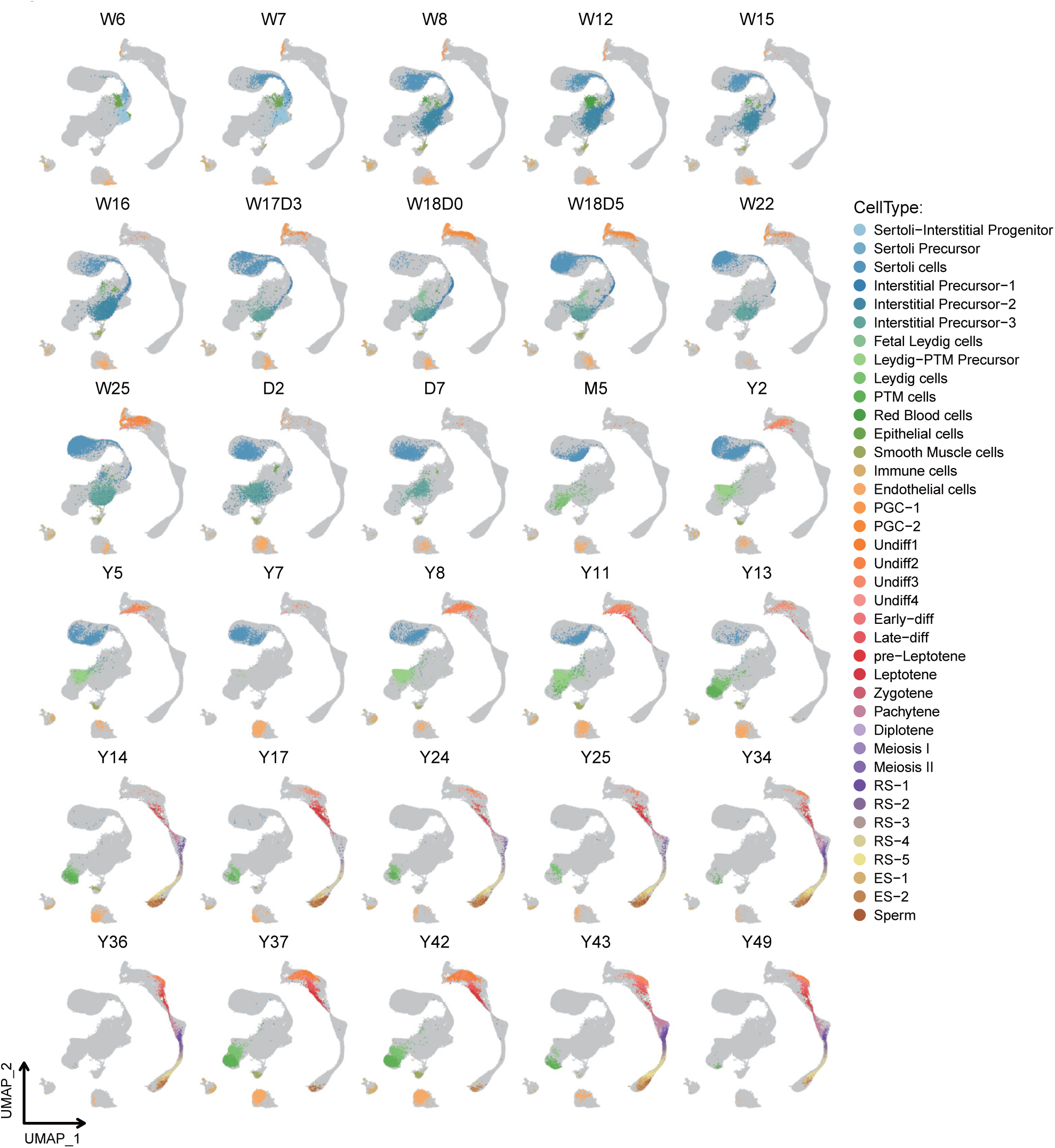
UMAP Partitioning based on age group, with cells colored by cell type.

**Figure S9.**
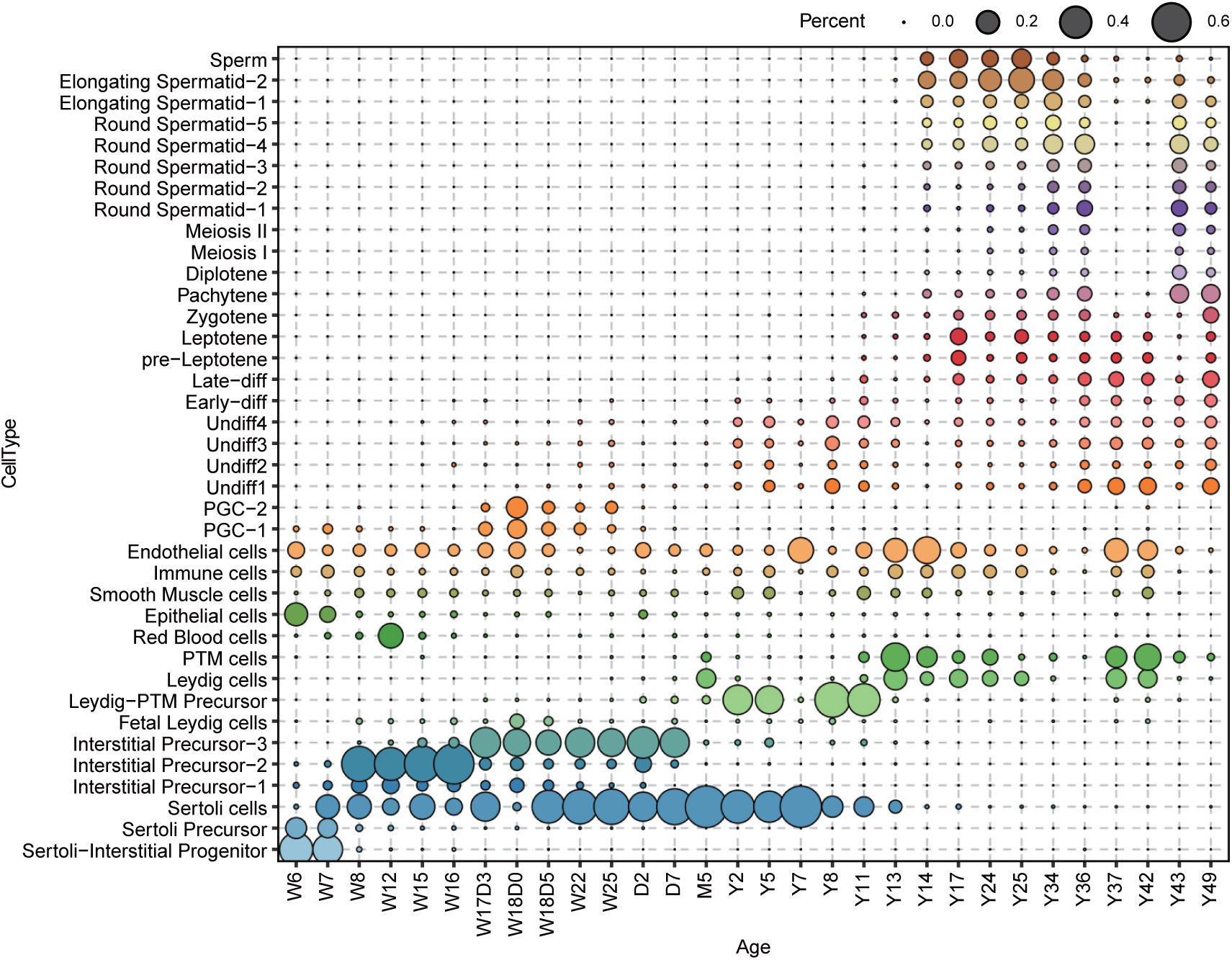
Statistical plot showing the distribution of each cell type of the “AllCells” cell set in each age group. The size of the dots is proportional to the percentage of cells.

## Tables

**Table S1 Detailed Information on data utilized by HumanTestisDB.**

**Table S2 Differentially expressed genes of undifferentiated Spermatogonia in Figure S4B.**

**Table S3 Key genes linked to functional biological events in spermiogenesis, as identified in the dynamic gene expressions of Figure S6C.**

